# Matrix-based methods versus Elo-ratings for assigning ordinal dominance ranks: comparisons in the Amboseli baboons

**DOI:** 10.64898/2026.11.21.768337

**Authors:** Jacob B. Gordon, David Jansen, Niki H. Learn, Jenny Tung, Elizabeth A. Archie, Susan C. Alberts

**Affiliations:** Duke University; University of Notre Dame; Princeton University; Max Planck Institute of Evolutionary Anthropology

## Abstract

Dominance ranks have profound effects on individual behavior, health, and fitness. Hence, establishing reliable and feasible methods for measuring individual dominance ranks is essential for researchers studying animal societies. Here we explain the process for measuring ranks followed by the Amboseli Baboon Research Project (ABRP), a long-term study of wild baboons in Kenya. We had two main goals. First, we document our matrix-based method for assigning ordinal dominance ranks to the baboons. Second, we compare our matrix-based method with another popular method that is frequently used in animal studies, the Elo rating method. We describe three aspects of the Elo rating method that can create inaccurate ordinal ranks in our system (and potentially others), at least in the short term. First, the Elo rating method constrains the pace at which an animal can rise through the ranks, relative to what we observe in baboon societies. Second, the Elo rating method assumes that absence of evidence is evidence of absence, i.e., it assumes that we have perfect knowledge of all interactions in the system. This assumption is inaccurate given constraints on how granular the observations in our study population can be. Third, the Elo rating method gives disproportional weight to unexpected interactions. We describe some of the consequences of each of these three aspects of the Elo rating method in our dataset on the Amboseli baboons. In the aggregate, we find that the matrix-based ranking method is more flexible, accurate, and easier to implement for our purposes.

## Introduction

Animals in many societies, especially primate societies, can be ranked according to their ability to win conflicts within their social group or community. Many methods have been proposed for assigning the “dominance ranks” that reflect these wins and losses. Methods fall into two broad categories (reviewed in de Vries 1998, Albers & de Vries 2001). Those in the first category (e.g., matrix-based, or “I and IS” methods; de Vries 1998) derive a ranking by optimizing features of the matrix of agonistic interactions as a whole. Those in the second category (e.g., Elo rating; Albers & de Vries 2001) score individuals on their success in winning contests, and derive a ranking by comparing individuals’ scores.

Each method has strengths that are well-suited to particular research goals, behavioral contexts, and data types. Many of these methods have been described and discussed ably and in detail elsewhere (e.g., de Vries 1998, Albers & de Vries 2001, Neumann et al. 2011, Newton-Fisher 2017); we do not review or assess most of them here. Instead, our goals are twofold. First, we document our matrix-based method for assigning ordinal dominance ranks to the baboons we study with the Amboseli Baboon Research Project (ABRP). Second, we compare our matrix-based method with another popular method that is frequently used in animal studies, the Elo rating method. We made this comparison in order to determine whether the Elo-based method produces dominance ranks that are comparable to the matrix-based method that we have used for many years (e.g., Hausfater 1975); the Elo-based method is attractive because it can be automated to a great degree, thus saving time and effort compared to the matrix-based method.

In the baboon population we study, individual dominance rank predicts a wide range of phenotypic outcomes, from life history and behavioral traits (e.g., Alberts et al. 2006, Gesquiere et al. 2018) to epigenetic aging and gene regulatory patterns (Anderson et al. 2021, Anderson et al. 2022). Individuals’ ranks change over time and our data are longitudinal. Consequently, to examine these relationships for any given set of animals during any given time period, we must assign ordinal ranks during specified time periods. Both matrix-based methods and Elo-based methods are potentially suitable for this type of analysis: both can produce ordinal dominance rankings for specific time periods that can be used as predictors of other traits. Matrix-based methods lend themselves easily to this approach, as they typically consolidate interactions over a specified time period to determine a single ranking for that period; a series of time periods (e.g., a series of months) can then be used as a time-varying predictor variable in mixed models. Elo ratings, too – which are designed to continuously track individual rank trajectories – can be used in such analyses by taking ‘snapshots’ of Elo scores and deriving ordinal dominance ranks from them.

Here we describe the use of both matrix-based and Elo-based ordinal rank assignments in our study population of baboons (*Papio cynocephalus* with admixture from *P. anubis*). We provide both general overviews of each method and descriptions of how we apply each method in our study population. We also provide a detailed comparison of dominance ranks produced by the two methods. We examine the correlation between the ranks themselves, and we provide comparisons of analyses using the two methods. In comparing the two methods, we identify several features of the Elo-based approach that can create inaccurate ranks in our study system. We provide specific examples of how the Elo rating method produces inaccurate ranks and end by concluding that the matrix-based ranking method is more flexible, accurate, and easier to implement for use in our study population.

## Matrix-based Ordinal Ranks

Each member in a baboon social group is assigned a unique numerical ordinal rank that represents its position in the group’s dominance hierarchy relative to all other same-sex members of the group for a particular month (i.e. ranks are tabulated for groups for every month). The ranks are determined by the outcome of decided agonistic interactions within dyads (see glossary below, and Alberts et al. 2020 for a description of decided agonistic interactions).

Briefly, sex-specific dominance rankings are calculated on a monthly basis for all females and all males (relative to other females and other males, respectively, in the same social group in the same month). We calculate sex-specific ranks because all subadult and adult males are much larger than all females, and they win fights over all females in dyadic contests. Additionally, work in this population and others indicates that the determinants and consequences of rank differ for males versus females. Thus, the hierarchy naturally sorts itself into distinct male and female hierarchies. However, we retain records of all agonistic interactions (including those between opposite-sex dyads) and we can create a whole-group matrix as well as sex-specific matrices. We are therefore able to examine cross-sex agonistic interactions in special cases (e.g., when we want to investigate the patterns exhibited by juvenile males as they mature and rise in rank over adult females). In the protocols described below, we only consider same-sex dominance rankings.

Ranks are calculated by generating an NxN matrix (where N is the number of individuals of a given sex in the social group). When we assign ranks to the Amboseli baboons, we include all individuals that were present in the group for at least a day during the month, but researchers may decide to assign ranks to just a subset of the individuals (for instance, only adult males who were present for at least half the month), as the method is flexible. The matrix contains symmetrical rows and columns, each corresponding to an individual animal’s identity. The cells of the matrix contain the number of times that the animal represented by a given row won an agonistic interaction against the animal represented by a given column in that month.

Generally, when researchers use a matrix-based approach in a species characterized by linear dominance hierarchies, several simple rules of thumb are followed. <u>First</u>, the relative ranking of any pair of individuals is considered stable from one time period to the next unless the behavioral data show evidence that the relative ranks of the two members of the dyad have changed (below, we provide details of what constitutes such evidence). <u>Second</u>, in assigning ordinal dominance ranks, the columns and rows of the matrix are ordered to minimize the number of wins that appear below the diagonal of the matrix. The resulting order of the columns (and rows) is the ordinal rank (e.g. 1, 2, 3, etc.) of the animals represented by those columns. <u>Third</u>, a given individual can and usually does lose to many other individuals that rank above it. Therefore, a given dominance matrix will typically have values in many cells above the diagonal (and comparatively fewer non-zero cells below the diagonal), indicating that multiple animals have lost in interactions with animals that rank higher than them in the hierarchy.

<u>Fourth</u>, researchers must decide whether to reorder the ranks when a “reversal” occurs in the matrix. We define a reversal as follows: (i) an individual appears to have only losses and no wins against an individual ranked below it, or (ii) both members of a dyad have at least one win against the other. In both of these cases, members of that dyad will have entries below the diagonal of the matrix and the researcher must decide whether these reversals constitute evidence that the rank order of the individuals has changed. Note that this use of “reversals” does not equate to “two individuals have reversed in their relative rank order”, rather it refers to a situation in which entries occur below the diagonal. In some cases, reversals (entries below the diagonal) will remain after the rank order has been finalized for a given time period, while in other cases reversals will be resolved because the ranks will be reordered.

As a general rule, we consider male dominance ranks to be more dynamic, i.e., to change more often and more rapidly, than female dominance ranks, and so we employ somewhat different rules for assigning male dominance ranks than for assigning female dominance ranks. For extensive details about how male ranks are determined, see the “Ranks” section of the behavioral and group movement data manager’s protocol (Gordon 2022) and for more details about how female ranks are determined, see the “Ranking” section of the demography and reproductive portion of the demographic and reproductive data manager’s protocol (Learn 2023).

There is a clear mathematical component to the process of assigning dominance rank, in that any agonistic reversal identifies a potential dominance rank change and having the smallest sum of reversals – i.e., the smallest number of entries below the diagonal of the matrix – is theoretically the most correct ranking. Indeed, often the total number of wins vs losses between two members of a dyad clearly indicates a rank change has occurred. However, occasionally the total number of wins vs losses is not sufficient to determine the outcome, e.g. when data from the dyad of interest are sparse or interactions with other individuals point to a different outcome. Which specific factors “matter”, and how much, varies according to the sex of the ranked individuals. These specifics are discussed below.

<u>First</u>, for each month we create a dominance matrix that includes every individual of a given sex who was in the group for at least one day during that month. Initially, they are each ranked using the order of the previous month’s ranks.

Second, we place males or females who are new to the group for that month in their respective hierarchy, using the following rules:

- Newly born infants are added at the bottom of the hierarchy appropriate for their sex. When multiple infants are added in the same month, they are ranked by age (older > younger).
- For males:

- Immigrant adults are placed below all other adults and above all subadults and juveniles.
- Immigrant subadults are ranked among the other subadults, according to their age estimate (older > younger; Note: All new immigrants are assigned age estimates by the field team at the end of the month in which they join).
- For females:

- Visiting adult and juvenile females are, unlike visiting males, extremely rare. Generally, visiting females are not ranked because they usually stay in the group only briefly and do not participate in enough agonistic encounters to provide reasonable guidance for placing them within the dominance hierarchy. Visitors present for less than two weeks will not be ranked. Longer-term visitors may be ranked on a case by case basis, with adults initially placed below the lowest-ranking adult female and juveniles placed by age amongst the juveniles still ranked below the lowest-ranking adult.

<u>Third</u>, the rank order is rearranged in order to minimize the number of agonistic interactions below the diagonal. In particular, for every “win” below the diagonal, we check the corresponding “loss” box above the diagonal.

- If the value in the cell above the diagonal is greater than the value in the corresponding cell below the diagonal (i.e., wins for the higher-ranked male exceed losses for that male, to the same interacting partner), or if the values in the two cells are equal, we do not record a change in rank between the two individuals in question.
- Otherwise, whichever cell has the higher value determines the overall winner in the dyad, i.e., determines who is higher ranking.

- Caveat for males: regardless of wins and losses, adult males who are only present in the group for a few days of a month (< 1 week) are ranked at the bottom of the adults.
- Caveat for females: with the exception of females floating between fission products during and soon after a fission, visiting females who are only present in the group for part of the month are generally not ranked.

<u>Fourth</u>, we consider cases in which we are likely to leave the existing hierarchy in place despite the presence of reversals (i.e., entries below the diagonal). In making decisions about when to re-order the rankings versus when to leave entries below the diagonal, we follow these rules:

- For adjacently ranked individuals in which a reversal occurs:

- In the case of males, a 1-0 win against the adjacent upper neighbor (i.e., the male ranked immediately above) in a single month is sufficient to indicate a change in rank, unless the upper neighbor won consistently for the 3-6 months prior to AND after the reversal. Adjacent males frequently flip back and forth from month to month; each flip is recorded as a change in rank for the month.
- In the case of females, a 1-0 win against the female ranked immediately above in a single month may or may not be sufficient to indicate a change in rank. Unlike with males it is very rare for pairs of females to frequently switch ranks. Female ranks tend to be more stable than male ranks so if a female is generally losing to the female ranked just above her in the adjacent months she will not be credited for a win that goes against this pattern; her rank is unchanged and stability is preserved. If, however, there are no other interactions between the pair for many months after this interaction, then a change in rank between this pair of females is recorded.
- For non-adjacently ranked individuals in which a reversal occurs:

- In the case of males, if a male wins only once over a male more than one rank above him (i.e. over a male that is higher ranked than his adjacent upper neighbor), don’t change his rank unless that win or a win over other males in between the two is confirmed in the next 1-3 months (or longer if data are scanty).

- One exception to this rule is when a male wins many times over a male more than one rank above him, e.g., 5-6 or more wins. In that case, give him credit for the rise, but watch both him and the male he won over carefully for the next few months to see if the loser is dropping precipitously or the winner is rising.
- Another exception is when a maturing male wins over a male several ranks above him, especially if the maturing male then disperses before he interacts with anyone else. In this situation, give the maturing male credit for the rank rise. In general, these young “rising” males get more credit for multiple “jumps” up the hierarchy than do older established males, especially if they are obviously on an upward trend anyway (see Hamilton & Bulger 1990).
- Sometimes a male wins over a male more than one rank above him several times in a month, but over no males in between. In order for a lower ranking male to get credit for rising in this situation, he cannot have any losses in that month to the males in between them, and either (1) he must win over a male in between in the next few months, (2) the higher ranking male must keep winning over the males in between during the same period that the lower ranking male is winning over him, or (3) the lower ranking male must be a maturing male as described above. Alternatively, for situations in which none of these things occur, and in which the higher ranking male loses to males in between in the following month, it may be that the higher ranking male is dropping in rank.
- In the case of females, when an adult female has a single win over a female more than one rank above her, a change in rank may be warranted, provided moving her up in rank would not cause new reversals with any of the females in between and the female over whom she has a win does not in turn have a win during the next several months.
- Whether the animals are adjacently ranked or not, in some cases a reversal is ‘contested’, i.e., a reversal occurs where the cross-diagonal box is not 0 (for instance, the lower ranking individual won over the higher ranking individual 4 times but lost 3 times), the following rules apply:

- In the case of males, contested reversals should not be counted as a change in rank unless a rank change is confirmed in the agonistic interactions data in the following several months.
- In the case of females, whether or not a rank change occurs depends upon: (i) the total number of wins for each female, both within the current month and going forward over the next several months; (ii) the timing of these wins; and (iii) any interactions the two females have with other females who rank between them during that time. Some additional circumstances should be considered:

- If the lower ranking female of the pair has more wins than losses within this window – considering all pairwise interactions with the opponent in question as well as females that may rank between them – then the lower ranking female is assigned a change in rank beginning at her first win, i.e., she rises in rank above the opponent.
- The lower ranking female in a contested reversal will be assigned a higher rank even if, in the current month, she has fewer wins than losses in one circumstance:
  - When at least one of the lower ranking female’s wins occurs later in the month than her opponent’s last win, and when she is clearly winning going forward over the next several months. If however, after adding in the new reversals, entries below the diagonal increase (i.e., she would have more losses than wins in her new, higher position), then no change in rank is recorded.
- However, the lower ranking female in a contested reversal will not be assigned a higher rank if doing so would result in more reversals overall due to three or more females having reversals with one another. In this case, whatever order results in the fewest agonistic interactions remaining below the diagonal should be used.
- Cases in which we do allow large ‘jumps’ from a lower to a higher rank occur in the following circumstances:

- In the case of males, place immigrant males over all males that they clearly win over in the first few months after immigration, even if there are multiple ‘jumps’ in these months, and especially if the data are scanty. In general a newly immigrant male (especially young and prime adults) will challenge the highest ranking males in the hierarchy without necessarily working his way up systematically from the bottom (Hamilton & Bulger 1990). This pattern also applies to maturing natal males who often make big jumps in rank during the attainment of adult rank status, as discussed above.
- In the case of females, adolescent females, especially those from higher ranking families, often make large ‘jumps’ as they move up the rank hierarchy toward their closest female relatives, ignoring many of the females in between. Since we know females usually attain rank near these close female relatives (Lea et al. 2014) we can anticipate the rank that these young females will eventually attain as they ascend through the hierarchy, and we can confidently allow large changes in rank even when we have observed only a small number of interactions. Sometimes a low-ranking adult female will have a small number of wins over an adolescent female during the same months in which the adolescent begins to accumulate wins over much higher ranking females. Provided the adolescent has further wins going forward and no losses to other intervening females, these losses should not prevent her from attaining a change in rank.
- Occasionally it is clear that a lower-ranking individual is not moving up *per se*, rather the higher-ranking individual is moving down, as evidenced by a large number of losses by that higher-ranking individual to multiple individuals over a period of several months. In such a case, contested or not, the individual who is losing rank is moved lower in the ranks without changing the order in which any other participants are ranked unless additional interactions amongst them indicate that a change in rank is warranted.

To sum up, when considering a change in rank it is always helpful to look at the 6 months or so before and after a rank change to get an idea of where a baboon is going and whether she or he confirms the wins/losses accumulated in a particular month. However, due to biologically relevant differences among individuals, particularly their age-class, confirmations are not always necessary to record a change in rank.

## Elo-based Ordinal Ranks

The Elo rating system is a method for calculating the relative skill levels of players in zero-sum games such as chess. The calculations are based on continuous updates that consider an individual’s interactions and their relative winning probabilities based on the current Elo scores (described below) of both winner (actor) and loser (actee). The Elo rating system has been widely used in animal behavior to assign both cardinal (continuous) and ordinal dominance rankings to individuals within social groups (e.g., Albers & de Vries 2001, Neumann et al. 2011). Here we briefly describe how Elo ratings might be used to obtain ordinal dominance rankings within baboon social groups, and then we consider some of the pros and cons of using Elo versus the matrix-based ordinal rankings that we described in the previous sections.

As with the matrix-based ordinal rankings, we construct separate rankings for male and female baboons. To assign ordinal rankings for a given month based on continuous Elo scores, we examine all the Elo scores at the end of that month and we order the individuals in the group according to the magnitude of their Elo scores, from the highest Elo score (highest ranking individual, assigned an ordinal score of 1) to lowest Elo score (lowest ranking individual, assigned an ordinal score of N where N is the number of individuals in the hierarchy).

An important point to consider is that the matrix-based method is designed to assign ordinal dominance ranks during a specific time period, which is represented by all the interactions that occurred during that time period and that are included in the matrix. In contrast, the Elo-based method involves continuous tracking of Elo scores (and thus relative ranks) over time. By creating monthly ordinal ranks using the Elo method we are taking snapshots of the Elo scores at forced time intervals.

In addition, in the Elo rating system, an individual’s Elo score can change only if the individual interacts. However, the scores of other individuals ranked around that individual can change, with the result that the individual’s relative ranking can change despite that individual having no interactions.

Specifically, Elo scores are based on the expected probability of an individual winning given its own current score and that of its opponent. In Elo scoring, each individual’s first Elo score (the ‘entry score’) is set to a predefined constant, typically a low value. Elo scores are then updated after each observed dominance interaction between two individuals; in each case, the winner receives a ‘winner’s bonus’, which increases their Elo score, and the loser pays a ‘loser’s tax’, which decreases their Elo score. The winner’s bonus and the loser’s tax are the same in magnitude, and depend on two quantities: (i) the predicted probability that the winner wins (based on the difference between the Elo scores of the winner and loser, prior to the encounter) and (ii) a predefined constant ‘k’.

Following the EloRating package in R (Neumann and Kulik 2019; see also Batchelder and Bershad 1979 and Albers and de Vries 2001) we generate Elo scores by first calculating each actor/actee’s probability of winning/losing an agonism, then adjusting their current Elo score relative to that probability. In doing so, we assume that the scaled difference in Elo scores prior to the interaction can be treated as a z-score, and therefore can be converted into a probability of winning based on the cumulative distribution function for the normal distribution.

If P_ij_ is the probability that individual i wins over individual j based on their current scores then,

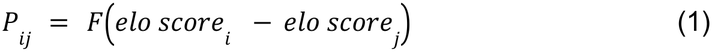

Here F is some form of a cumulative distribution function such that

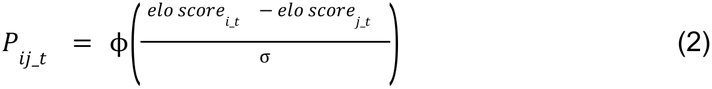

Where **φ** is a normal distribution and σ is an arbitrary scaling constant (here 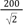).

The new elo scores for any individual i after an interaction with any individual j can then be calculated as a function of the winning probability for i against j, the old scores for both i and j, and the constant ‘k’, a scaling value.

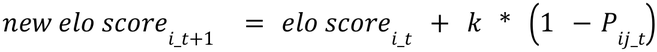

And similarly:

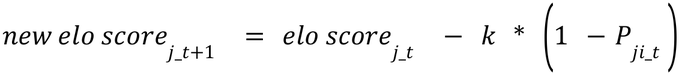

The k-value determines how much each interaction affects the Elo score. The k-value can be thought of as determining the magnitude of the ‘winner’s bonus’ and the ‘loser’s tax’ in an agonistic interaction (Franz et al. 2015). With larger k-values, the effect of a single interaction is bigger in changing the Elo score (and thus the relative ranking) of a given individual. We initially set the value of k at 100. We also varied the value of k in a series of exploratory analyses, and found that varying k did not greatly affect the correlations between Elo-based ordinal ranks and the matrix-based ranks (see section entitled ‘The “k” constant’ below). Hereafter we report results using k=100.

The code we used was based on the EloRating R package. We adapted the code to more closely match the ABRP data structure and baboon rank-related behavior, and to facilitate the extraction of monthly ordinal ranks. All of this code is available at https://github.com/ArchieLab/ABRP-Elo-based-ranks/.

## Comparing Elo-based and Matrix-based Ranks

Naturally, there are discrepancies between Elo-based ordinal ranks and matrix-based ordinal ranks. In many cases these discrepancies are small and inconsequential; in the aggregate, ranks produced by the two systems are highly correlated (Figure 1).

**Figure 1.**
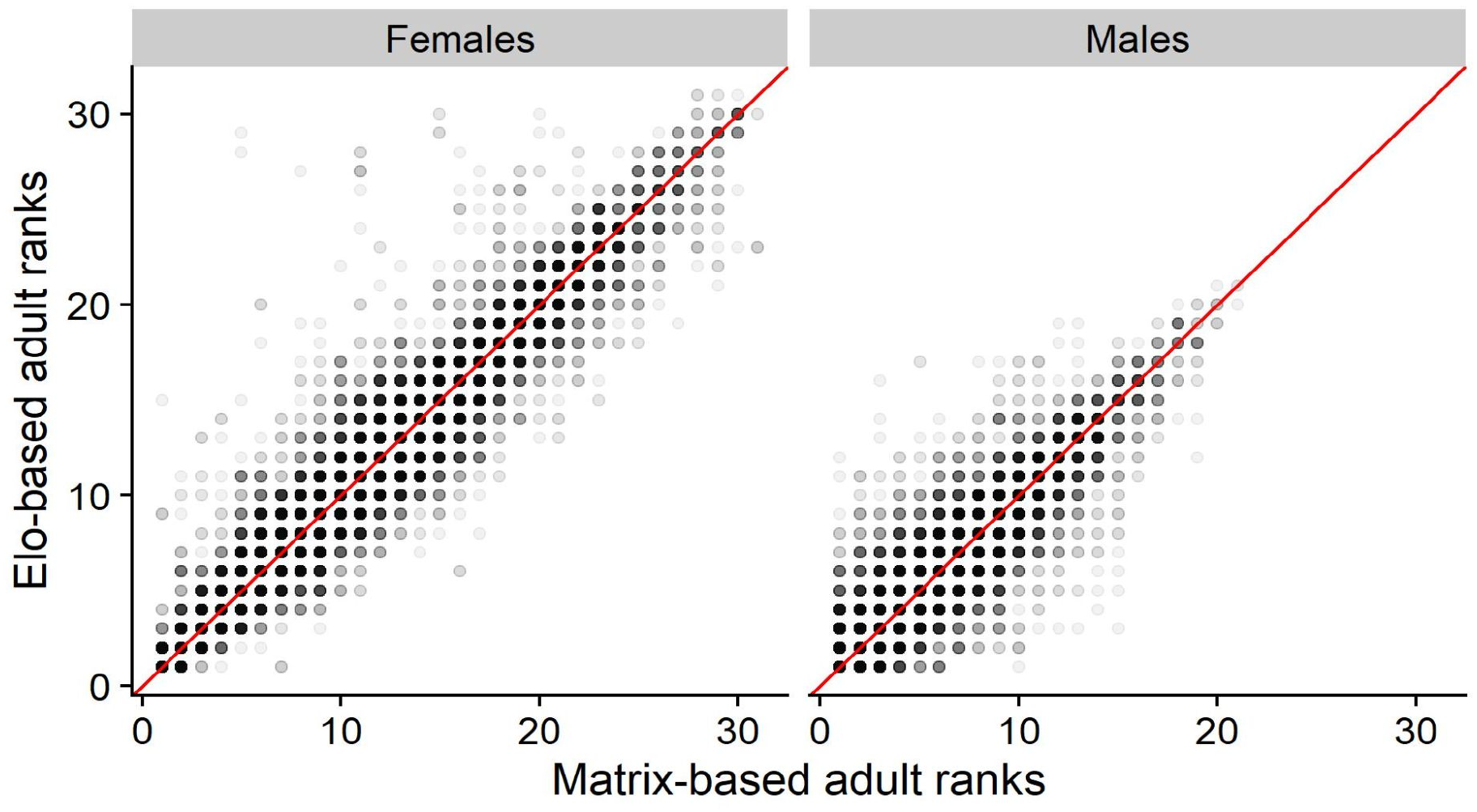
Relationships between matrix-based ordinal ranks (x-axis) and Elo-based ordinal ranks (y-axis) for female (left) and male (right) baboons in the Amboseli study system. The correlations between the two methods are high: for males, r=0.913, n=48,000 monthly ranks; for females, r=0.976, n=57,885 monthly ranks. Data were collected between 1980 and 2022 from, respectively, 941 and 777 unique males and females.

For 6.7% of monthly ranks for males and 13.6% of monthly ranks females, ordinal dominance ranks assigned by the Elo method and the matrix method differ by more than 2 rank positions, and 3.24% and 8.42% differ by more than 4 rank positions.

Furthermore, Elo-based rank assignments and matrix-based rank assignments produce virtually identical results in many analyses that involve rank as a predictor of physiological, behavioral, or life history outcomes. For instance, the two methods are nearly indistinguishable when using male dominance rank to predict mating success (Figure 2).

**Figure 2.**
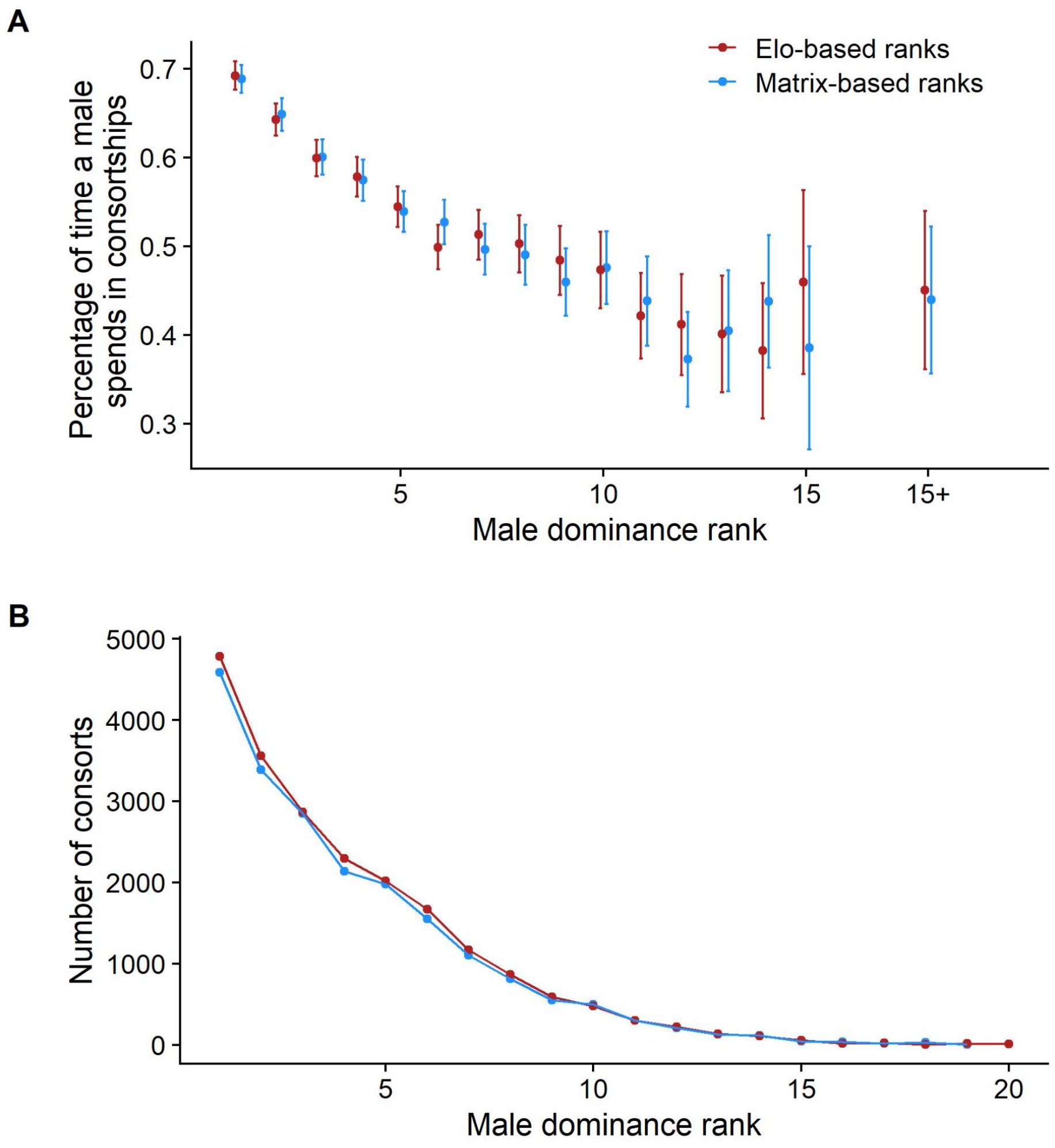
The relationship between male ordinal dominance rank and A. the percentage of time a male spends in consortships (mate-guarding episodes) and B. the number of consortships a male obtains. Here, matrix-based methods (blue) and Elo-based methods (red) of assigning ordinal dominance rank produce virtually identical results (A. r = 0.95; p.value < 0.001 and B. r = 0.99; p.value < 0.001).

The same is true when we use both methods to predict male hormone concentrations (as in Gesquiere et al. 2011), male paternity success (Alberts et al. 2006), female interbirth intervals and their component parts (as in Gesquiere et al. 2018), wound healing rates in males (as in Archie et al. 2014) and, in fact, almost every other analysis for which we have now compared both methods (Figure 3).

**Figure 3.**
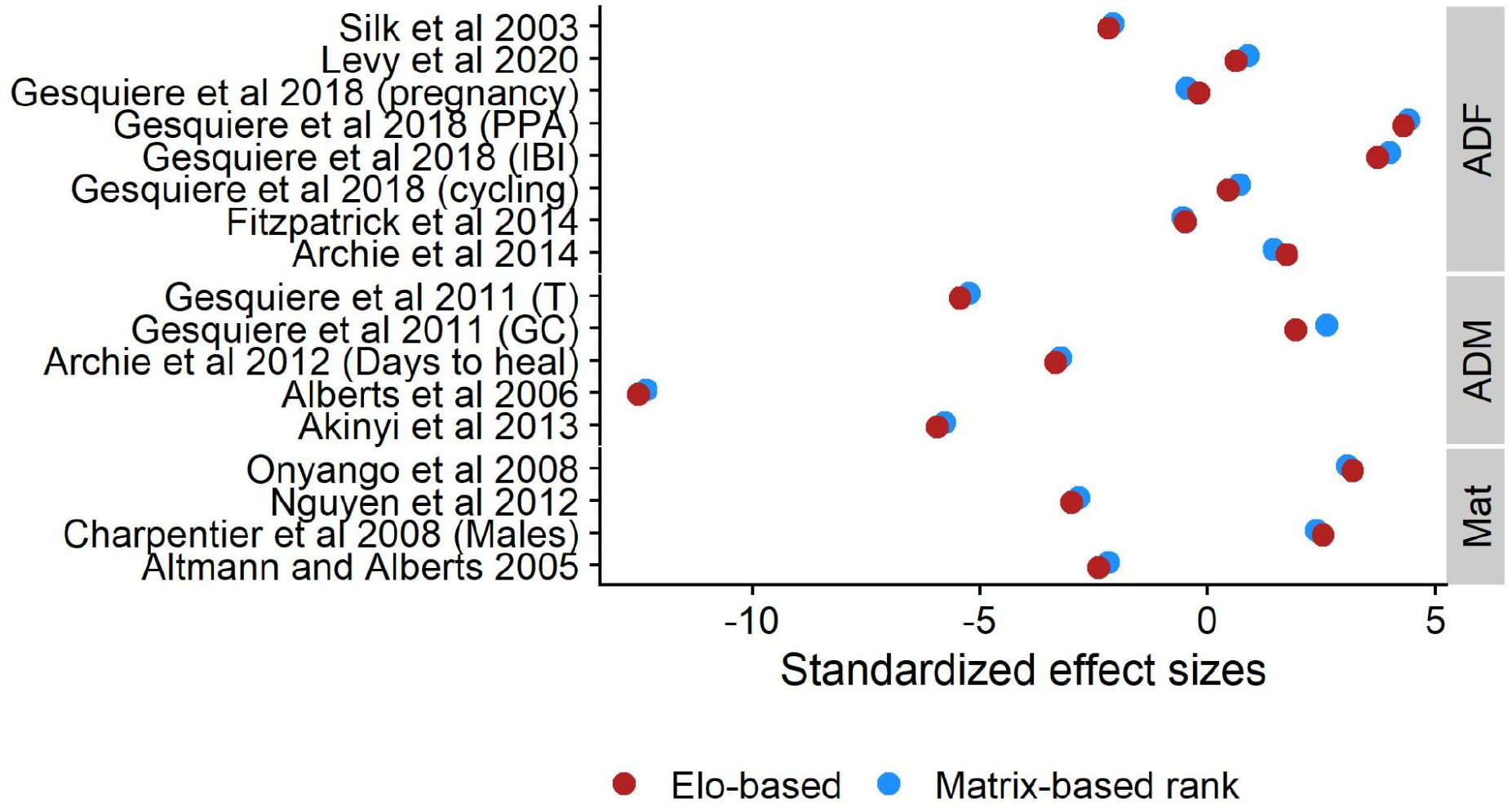
The standardized effect sizes of rank related variables in statistical models from a set of ABRP papers. The y-axis represents a list of published papers that included dominance rank as a predictor of a behavioral or life history outcome (see references). For each of the original analyses, we compared models that used matrix-based ordinal dominance ranks with models that used Elo-based ordinal ranks and extracted the estimated effect sizes for the rank measures. In nearly all cases, the standardized effect sizes for the effect (mean effect/std.error of the effect) of rank were extremely similar for Elo-based (red) and matrix-based (blue) ordinal ranks (r = 0.99; p.value < 0.001).

However, we argue that the reasons for these discrepancies are important for researchers to consider. In the following sections, we discuss those reasons. We do not claim that one system is correct and the other is incorrect. Rather, this discussion is intended to show that the Elo-based method is not a simple replacement for the matrix-based method. For example, we argue that the Elo-based method fails to account for some important biological and life history features of baboons, as well as some methodological aspects of data collection. We focus on three problems that arise from these failures.

### Problem 1: The Rate at which Individuals Rise and Fall in Rank

In general, to rise in rank an individual must win over higher-ranked individuals more than they lose to them. Ideally, if an individual rises in rank, for instance from rank #7 to #3, the data will include evidence that the individual who is rising can win over #4 - #6 as well as over #3. This is true in any ranking system. However, in baboon societies, some individuals – e.g. adolescent females or subadult males ascending to adulthood, or new immigrant males finding their place in the hierarchy – really do “jump” upward in rank without defeating everyone ranked between them. Different ranking systems use different criteria to decide when and whether to attribute rank rise to the ‘jumping’ individual.

In our system of assigning matrix-based ordinal ranks, we take a “big picture” perspective on such putative rank rises (or falls) by taking into consideration the maturational or immigration context of the individual’s agonistic interactions, and by examining the preceding and succeeding months, to get an idea of an individual’s position and trajectory in the group.

Looking at the “big picture” in this way might justify a change in rank (or lack thereof) that may not be explained otherwise. In contrast, the Elo rating system (i) does not take into consideration the differences between animal societies in how rank changes occur (for instance, it does not easily take into account special circumstances such as immigration and maturation) and (ii) importantly, it does not consider future events at all. As a consequence, individuals tend to rise or fall in Elo-based ordinal rank more slowly than they do in the matrix-based ranks.

This difference is clearly shown in the female ranks, particularly for adolescent females, in Yoda’s group in 2021. Most of the adolescent females eventually attain the same or similar ranks using either the matrix-based method or the Elo-based method, but they arrive at those ranks much more slowly under the Elo rating system than with our matrix-based methods, as shown for the four adolescent females (with IDs of RIJ, RIQ, YEE and YUY) shown in Figure 4.

**Figure 4.**
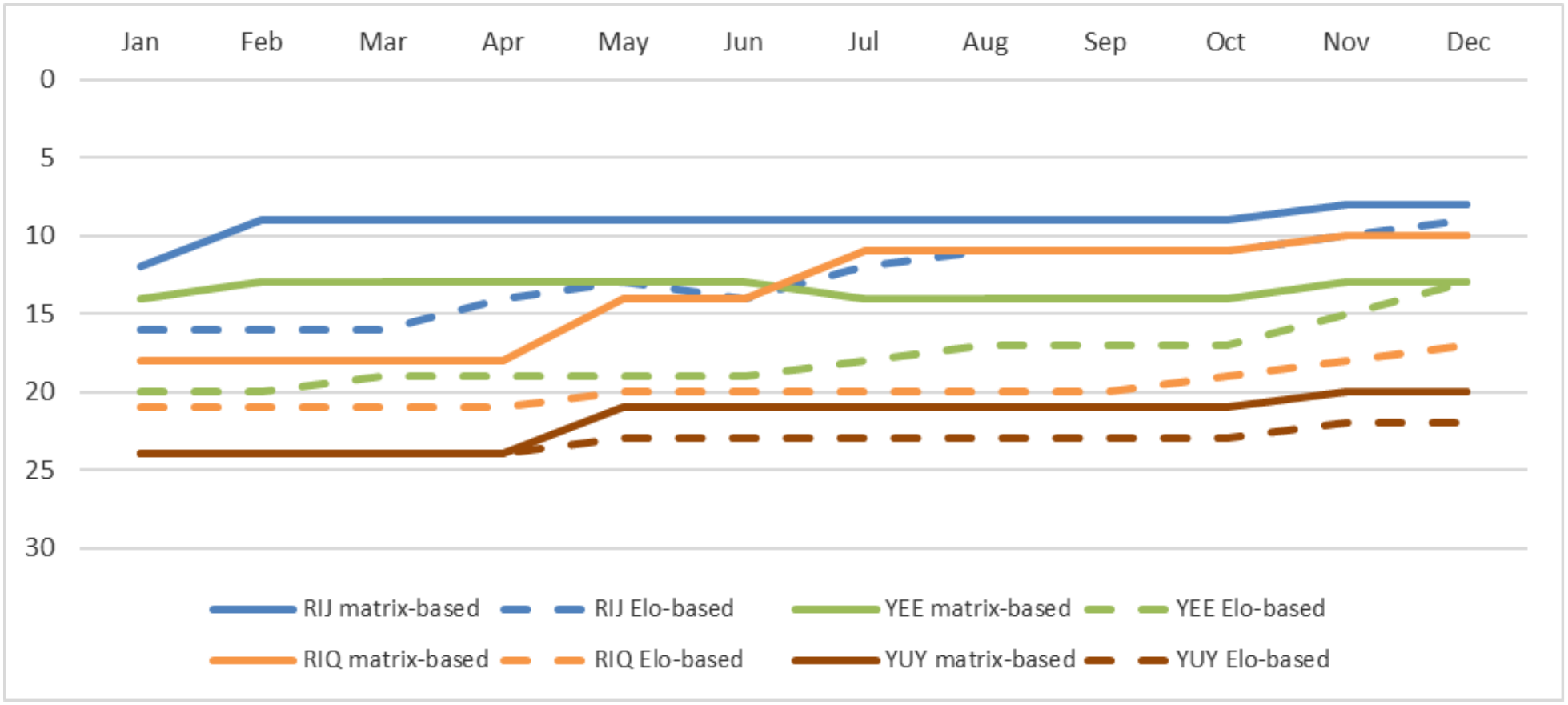
Graphical representation of four adolescent females rising through the female dominance ranks during maturation. The y-axis is ordinal dominance rank (higher ranks at top of axis, lower ranks at bottom) and the x-axis is month during the year 2021. When a female begins to rise rapidly through the ranks during maturation, such as with these four adolescent females in Yoda’s group over the course of 2021, the Elo-based method (dashed lines) is much slower to alter the rank hierarchy following their wins than is the matrix-based method (solid lines).

This type of discrepancy is also evident for male baboons in numerous cases. For example, in Yoda’s group, August 2021, recent immigrant TOI had several wins that secured his place as the #1 male in the group that month in our matrix-based assignments of ordinal ranks. However, in the Elo-based ranking system, TOI does not rise above several of the males that he clearly defeated (UPE, KOM). At the time of TOI’s immigration into the group, TOI was assigned an Elo score that placed him above all sub-adult males and below all adult males. At this point, the score difference between TOI and the top-ranked male UPE was approximately 1500 Elo points. This score difference gives TOI a low probability of winning against UPE, so when he does win against UPE he gets a relatively large ‘reward’, i.e., a large increase in his Elo score (see equations 1 - 3, above). However, with “k” set at 100 his maximum rise in score would be 100 for each win against UPE, while UPE’s score would simultaneously be reduced by 100 as a result of that loss. Even if there were no interactions with other individuals to affect their scores, TOI would need to win at least 7 times to surpass UPE, because as their scores converge, the “bonus” for winning decreases. This example illustrates an important point about the Elo-based method: it makes assumptions about inter-individual differences that have important implications for how we interpret an animal’s behavior.

As another example, in Hokey’s group in July 2021, subadult male HOT had several wins over (and no losses to) adult males. According to the rules of our matrix-based rank assignment, he rose above them and attained his “adult” rank for the first time in July 2021. The attainment of adult rank is an important developmental milestone in male baboons, and we assess its timing with some care (Alberts and Altmann 1995). However, our Elo-based ranking system keeps HOT ranked below all adult males for that month because at the start of the month his Elo score is much lower than the scores of other males ranked above him. His score does increase quickly due to ‘unexpected’ wins, but he doesn’t achieve a sufficiently high Elo score to surpass any adult-ranked males until August. In other words, even if an individual consistently wins conflicts against a higher-ranking individual, the Elo method does not allow the winning individual to rank above the losing individual until the winner accumulates enough points in the Elo method to surpass the losing individual’s Elo score. If the initial score difference is large it may involve multiple wins and therefore considerable time to overcome this Elo score deficit. In other words, the Elo rating method lends itself to more conservatism in rank hierarchy dynamics than we believe to be the case, especially for key age-sex classes.

#### The k constant

In response to this concern, one might ask about changing the k parameter, which is set to a constant value by the researcher. When k is set to a small value (relative to entry score, which is also set to a constant value by the researcher), single outcomes generally have only a small impact on changes in Elo scores. Small values of k thus assume that single wins and losses are not very predictive of future wins and losses. When k is set to larger values, single outcomes will have larger effects on changes in Elo scores, which implies that single wins and losses should be more predictive of future wins and losses. If the ranks of individual animals aren’t rising/falling rapidly enough in response to their interactions, doesn’t this suggest that our Elo algorithm should use a larger k, so that each individual interaction has a greater effect on an individual’s Elo score?

This problem is not solvable by changing the value of k for the dataset as a whole, because the k parameter affects the ‘winner’s bonus’ and the ‘loser’s tax’ for <u>all</u> individuals, not only those who may be experiencing rapid rank rises of the type described above. In other words, using a higher k constant generally means that we simply trade one problem (i.e., that rank changes need to be more rapid for some individuals) for a new one by creating a population in which rank changes overall are very (and perhaps overly) dynamic.

Another potential approach involves relaxing the constraints of the Elo-rating methods, for instance by allowing k to vary across individuals or across different types of interactions (e.g., Franz et al. 2015, Newton-Fisher 2017), or by allowing the individual’s entry score (i.e., initial Elo score at entry into the hierarchy) to vary across individuals (e.g., Foerster et al. 2016, Goffe et al. 2018, Feldblum et al. 2021). These methods allow the researcher to relax the assumptions that entry score and/or k are constant across individuals; allowing these parameters to vary depending on other variables – such as individual attributes or aggression intensity – helps correct rank assignment problems associated with too-slow or too-fast rank changes. However, these approaches are coding-intensive and time-consuming, requiring effort and careful consideration during the k-estimation phase. Furthermore, it is not clear how easily they can solve the problems that we describe here and below. These shortcomings make this approach somewhat impractical for broad use and application. Furthermore, while this approach is potentially effective for Problem 1, it does not address Problem 2 or Problem 3, which are described below.

#### Consequences of problem 1

Why does it matter if individuals don’t attain high rank as quickly in the Elo system as they do in the matrix-based rank assignments? In baboons, the consequences differ by sex.

##### Consequences for Females

The slower rank attainment of adolescent females in the Elo-based method can affect the ranks of the adult females in her path. Suppose there is an adolescent female who is on the rise, and she wins over an adult female who is ranked below her according to matrix-based assignment but above her according to Elo-based assignments (because attaining rank according to Elo-based assignments takes longer). When a lower-ranked individual defeats a higher-ranked one, the magnitude of the change in the Elo score for both individuals is relatively large. As a consequence, the Elo score of the defeated female may fall enough to bring her below other adults ranked near her, even if she never loses an interaction with any of them. In contrast, if the Elo-based system were capable of recognizing that the adolescent female had already risen above the defeated female, the magnitude of the change in the defeated female’s Elo score would be relatively small, and would be much less likely to impact her rank relative to other adults ranked near her.

For example, in Yoda’s group in March 2021, adult female EMC lost to adolescent female RIJ. In the matrix-based ranks, RIJ was already ranked higher than EMC at the start of the year, and was ranked well above her by February 2021 (Figure 5). In contrast, the Elo-based rank assignment for RIJ placed her three rank positions below EMC in March 2021. The negative impact on EMC’s Elo-based rank of her loss to RIJ in March 2021 was large enough that EMC’s Elo-based rank sank below that of YOD, the female directly below her. Importantly, YOD had never won over EMC, and EMC to that point had 12 wins over YOD under her belt, the most recent only three months earlier in December 2020. EMC went on to win over YOD again in June 2021 and then again in January 2022, leaving no doubt that YOD should not be ranked above EMC. Nevertheless, in Elo, YOD remains above EMC through June, not flipping back below EMC until July 2021.

**Figure 5.**
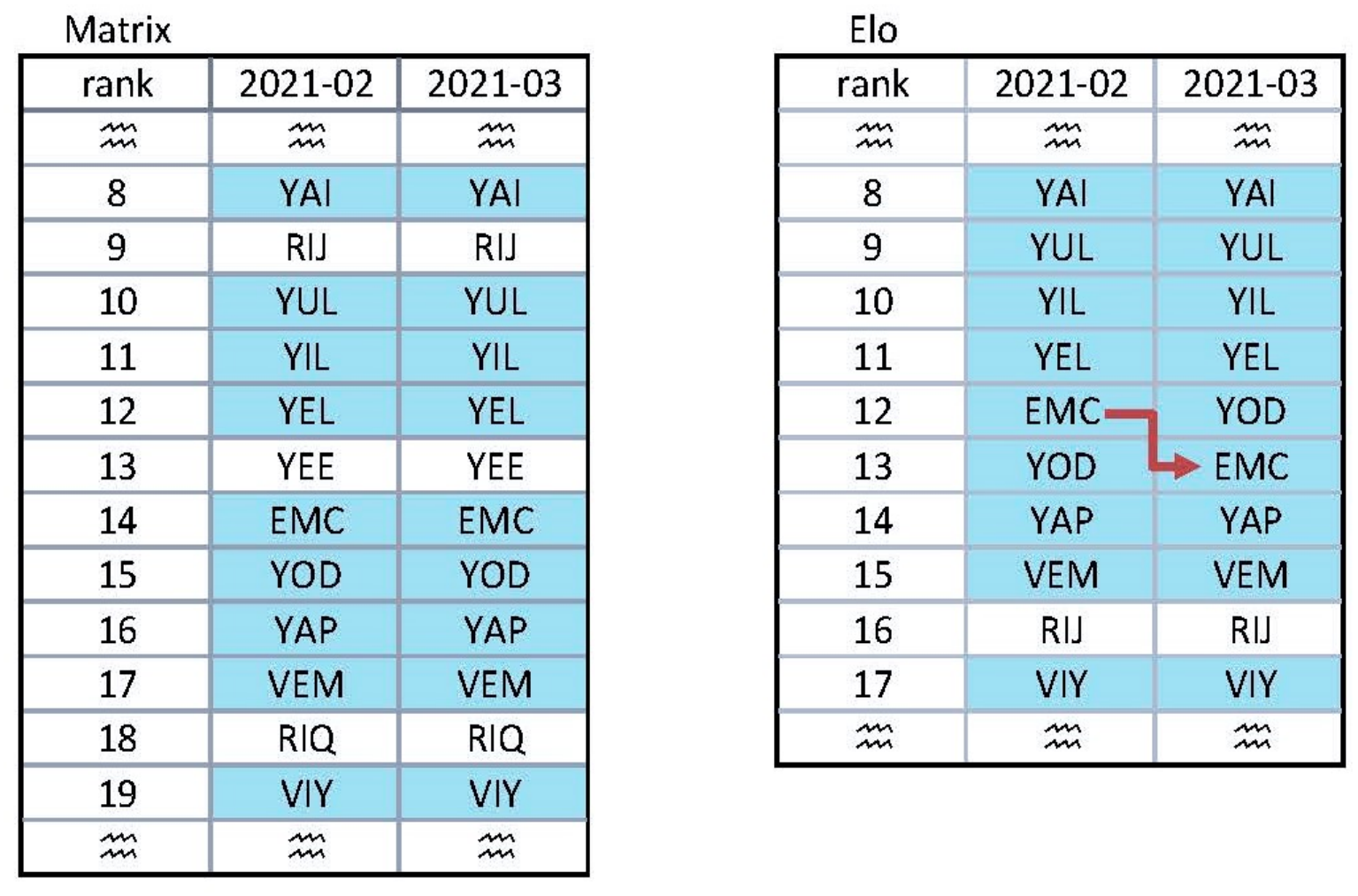
Excerpt from the female rank realignment sheet for Yoda’s group in February to March 2021. In March RIJ has a win over EMC. In the matrix-based ranks adolescent RIJ is #9, well above EMC at #14 and thus no reversal actually exists. No change in ranks occurs for either female in March. But in the Elo-based ranks RIJ is still only #16, three ranks below EMC. Since Elo does not expect RIJ to beat EMC, there is a negative impact on EMC’s Elo score that results in her sinking one rank below YOD, the adult female ranked just below her, despite the fact that YOD has never had a win over EMC. Mature females are highlighted in teal. The unhighlighted females shown are all adolescents who ascend in rank more rapidly in the matrix-based method than in the Elo-based method.

A detailed comparison of 2021 female ranks using the matrix-based method versus using the Elo-based method revealed many examples of females not changing ranks quickly enough in response to events, or changing ranks erroneously in response to interactions between other females around them. The result is the appearance of greater instability in the rank hierarchy with the Elo-based method than with the matrix-based method, with some females repeatedly switching rank positions due to small changes in their Elo scores. In other words, the Elo-based method paradoxically produced both slower rank rises for some individual females, and far more changes in rank across the year, than the matrix-based assignments.

###### Consequences for Males

We define a male baboon as an “adult” when he can consistently win fights with other adult males (see discussions in Alberts & Altmann 1995, Alberts et al. 2003, Alberts et al. 2006). This life history milestone has important functional consequences: subadult males perform virtually no mate guarding of fertile females and father few or no offspring (Hamilton & Bulger 1990, Packer 1979, Alberts & Altmann 1995, Alberts et al. 2003, Alberts et al. 2006). In contrast, once a male has attained a dominance rank among other adult males, he typically begins mate-guarding immediately, often within days or weeks. This milestone – the age at which adult reproduction begins – is of central importance for understanding the evolution of primate life histories.

The attainment of adult rank is often fairly sudden and rapid: one month, a male is clearly ranked below all the adults, then in the next month he has ascended to a new plane, fighting and winning against adult males that he previously consistently lost to (Hamilton & Bulger 1990, Alberts and Altmann 1995, Alberts et al. 2003). The slow attainment of adult dominance rank that occurs in the Elo-based system does not match this biological reality. If Elo-based ranks were used to assess when a male first attained adulthood, we would often be overestimating the age at which males attain this key life history milestone.

For example, three subadult males attained adult rank in 2021 according to our matrix-based ranking system: GON and HOT in Hokey’s group and ELK in Acacia’s Group. In the case of GON, our matrix-based ordinal ranks place him as an adult in November 2021, while in our Elo-based system he never scored higher than any of the adults before he emigrated out of the group in March 2022. In the cases of ELK and HOT, the discrepancy was not as great: their Elo scores rose above some of the adults one month later than the rank attainment date determined in our matrix-based ranks. However, overall these discrepancies would be expected to produce systematic bias towards later ages of maturation.

### Problem 2: Absence of Evidence, Evidence of Absence

Matrix-based rank assignments and Elo-based rank assignments make contrasting assumptions about the nature of the agonism data. In the matrix-based approach, we recognize that the agonism data are only a sample of all the agonistic interactions that occurred in the group that month. In contrast, the Elo-based approach implicitly assumes that the agonistic interactions in the dataset include all agonisms that actually occurred. That is a reasonable assumption for a chess or sports tournament, but not for a population of wild animals. Because of this assumption, Elo interprets an absence of evidence (i.e., an absence of interactions) as evidence of absence (i.e., as evidence that such interactions did not occur).

#### Consequences of Problem 2

Problem 2 has at least two important consequences. First, in the Elo-based ranking system, an individual can fall in rank by having few or no interactions. Suppose an individual ranked #3 in the group has few interactions with others in the group, while #4 wins repeatedly over #5 and #6. Individual #4’s Elo score will increase, while that of #3 will change little or not at all. This could result in individual #4 having a higher Elo score than #3 and therefore a higher ordinal rank, despite an absence of evidence to suggest that #4 is actually dominant over #3.

This situation occurred among the adult females in Europa’s group in February 2021 (Figure 6). Before this month, VES had been ranked #6 over TEL, #7. In the whole month, VES had just two agonistic interactions: a loss to #2 VOM and a loss to #5 TIB. These two losses decreased VES’s Elo score, but only minimally, as VOM and TIB were already ranked above her. TEL (#7) had many more losses to higher-ranked females but also had three wins over females that were ranked lower than *both* TEL and VES: TEL won twice over VEJ (#8) and once over USA (#9). Those wins increased TEL’s Elo score. With TEL’s score increasing and VES’s score decreasing slightly, Elo ratings placed TEL above VES this month, in the absence of any evidence that TEL was able to win a conflict with VES or any females higher-ranking than VES.

**Figure 6.**
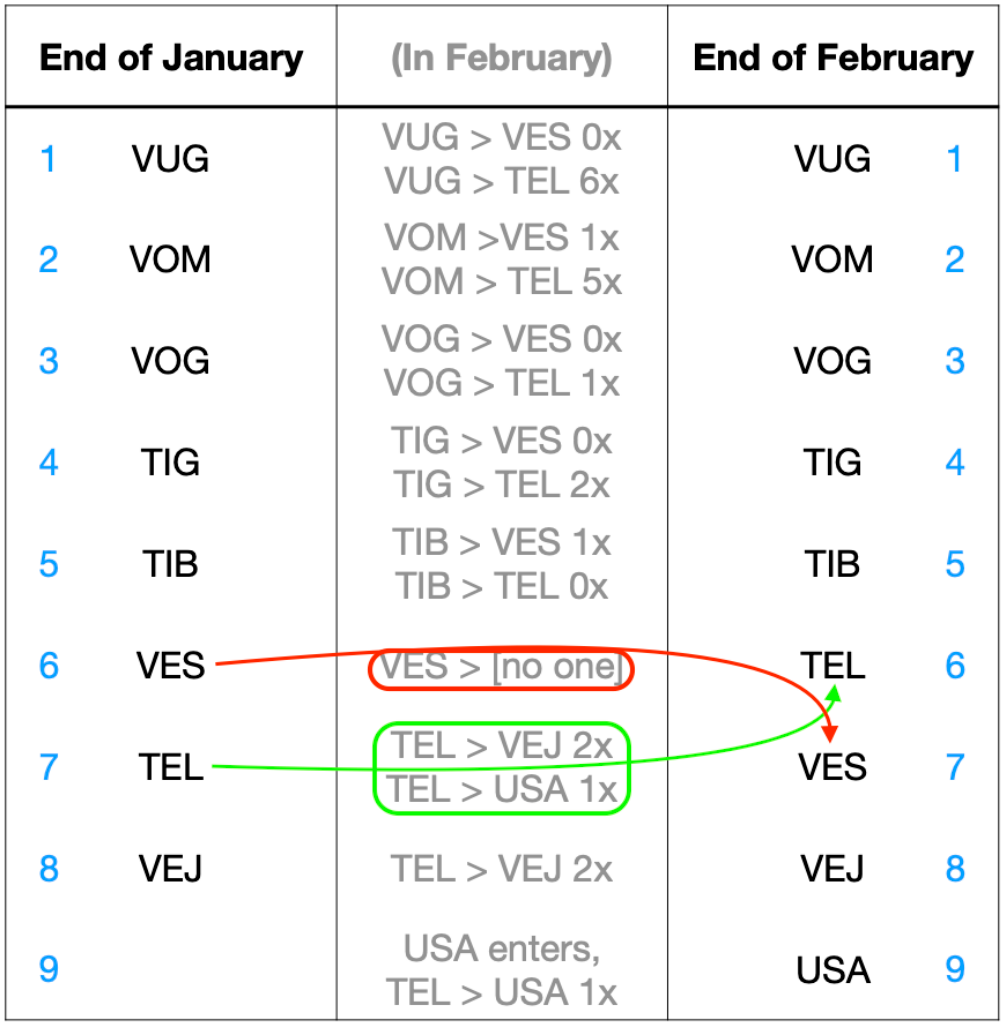
Excerpt of adult female ranks in Europa’s group in February 2021, including agonisms involving VES and TEL. Ordinal ranks shown in blue. VES had no wins in the entire month and a few losses to higher-ranked females, causing her Elo score to fall (red arrow). TEL had several wins over other lower-ranked females, causing her Elo score to rise (green arrow). This causes TEL to rise above VES, despite a total absence of interactions between them and despite the fact that all TEL’s wins were against females that were lower-ranking than both TEL and VES.

The symmetrical case is that an individual B can rise in rank above individual A if individual A experiences several losses to higher-ranked individuals, even in the absence of direct evidence that B can win over A. Suppose a high-ranking individual, #2 in the group, has several wins over #3 and #4 in a month, with the most wins against their nearest rival, #3. This would, in general, lower the Elo scores for both #3 and #4. Meanwhile, suppose that most of #4’s agonistic interactions are wins against lower-ranked individuals, with few or no interactions between #3 and #4. Between the aforementioned losses to #2 and these wins against lower-ranked individuals, the net impact on #4’s Elo score would likely be minimal. In contrast, if #3 loses multiple times to #2, then #3’s Elo score might fall enough for #4 to rise above her in rank. Again, this rank reversal could occur despite an absence of evidence to suggest that #4 is actually dominant over #3.

This unjustified rank change happened between VLA and VAQ, two of the adult males in Acacia’s group, in May 2021 (Figure 7). In that month, a new male (REY) entered the group and won five times over VLA. Just below VLA is VAQ, who didn’t interact with VLA at all that month, and always lost interactions with VLA up to that time. In our Elo-based rank system, VLA’s five losses to REY, who was an immigrant male undergoing a rapid rank rise (as per Hamilton & Bulger 1990), cause VLA’s Elo score to drop so much that he dropped below VAQ in ordinal rank, in spite of a complete absence of evidence that VAQ could win a conflict with VLA.

**Figure 7.**
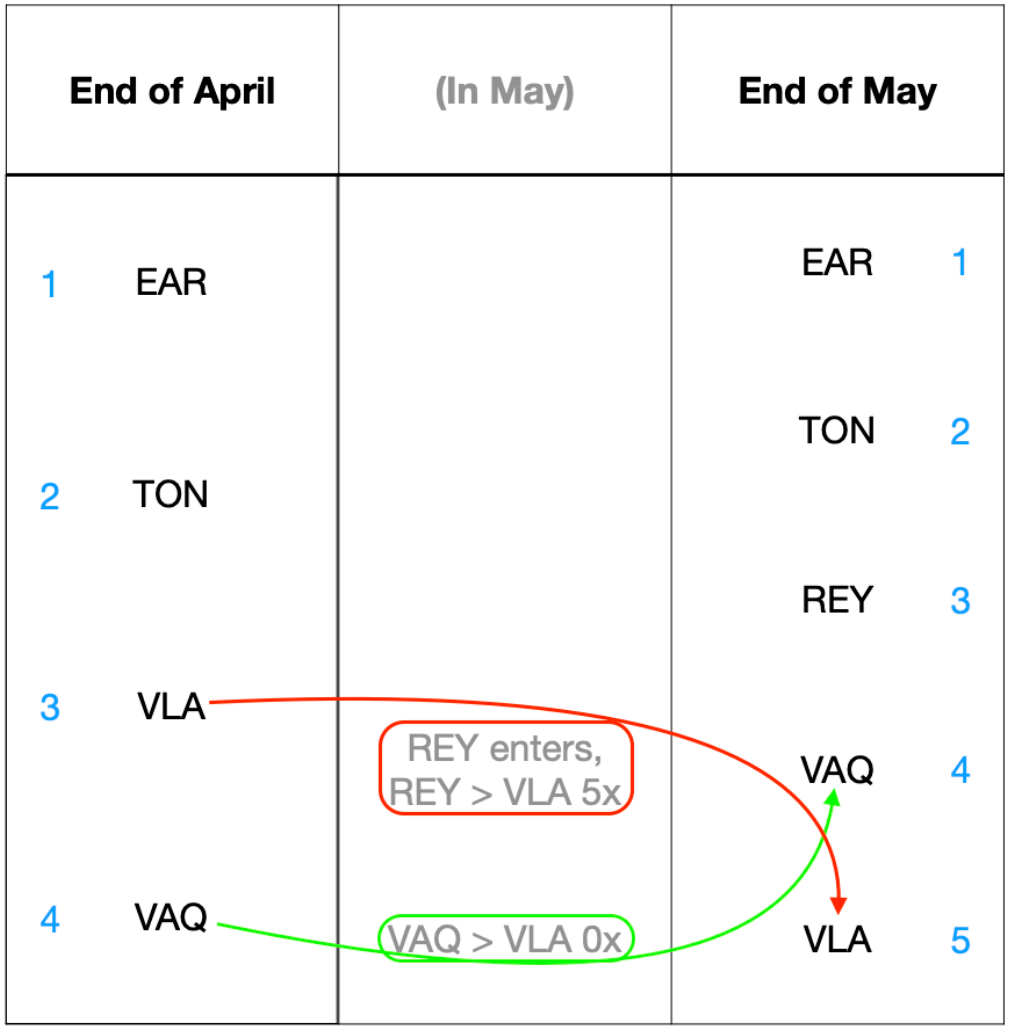
Excerpt of male ranks in Acacia’s group in May 2021, including agonisms involving REY, VLA, and VAQ. Ordinal ranks shown in blue. REY immigrated into the group this month, and had 5 unreversed wins over VLA. VAQ has no interactions with VLA, but his rank rises above VLA’s because of VLA’s five losses to REY.

### Problem 3. Unexpected interactions have a disproportionate effect on Elo scores

In a chess or tennis tournament, each competitive interaction is relatively protracted, and each pair of contestants will interact with each other relatively few times over the course of the particular tournament or the annual tournament cycle, or even across years. Most importantly, all interactions are equally significant (none are discounted) and each interaction is scored with no errors.

In contrast, in animal societies, competitive interactions between pairs of animals will be repeated over many months, years, or even over lifetimes, and not all competitive interactions are likely to be equally significant to the contestants. In addition, agonistic interactions are of relatively brief duration, datasets may be much larger (i.e., 100s or 1,000s of interactions), and observers may occasionally make an error in recording an interaction. Together, these features of animal societies and of the observers that collect data on them probably cause unexpected interactions to appear in the dataset somewhat more frequently than in sports rankings.

Importantly, single unexpected interactions have a disproportionate effect on Elo scores, as might make sense in sports rankings but perhaps less so in a very large dataset on animal conflicts. In an Elo-based ranking system, the existence and timing of highly improbable interactions (whether they are accurate representations of an unexpected event or are the result of observer error in recording the interaction) will have a disproportionate effect on the Elo scores for that month, and hence on the ordinal ranks derived from them. For instance, a given male can have many expected interactions (i.e., wins against lower-ranking animals or losses to higher-ranking ones), but as they are expected (have a high probability) they will have a limited effect on his Elo score. However, unexpected interactions (with a low probability, i.e. wins against higher-ranking animals or losses to lower-ranking ones) have a much larger effect on his Elo score. This will disproportionately affect his monthly ordinal rank, especially if it happens at the end of the month, as it may take many interactions to overcome this rise/drop in Elo score.

In contrast, our matrix-based method takes the opposite approach – a very unexpected event is scrutinized closely for evidence that it represents a real rank change. That is, in the face of an unexpected event, we examine previous months and future months for evidence that this rank reversal represents a probable change. This problem is related to, but somewhat distinct from, the “absence of evidence” problem discussed above. Our ability to consider the future as well as the past in our matrix-based rank assignments helps us to identify which unexpected interactions are highly salient for an animal’s rank in the group and which are not.

## Conclusion

The Elo-based ranking method and matrix-based ranking methods both have advantages when applied to animal societies. The Elo-based ranking method is an excellent method of assessing the overall winning success of an individual in social contests. In contrast, the matrix-based method, we argue, more carefully considers the biology of the organism (especially heterogeneity in rank dynamics due to interindividual differences in age, sex, or other characteristics) as well as individuals’ positions relative to others. Importantly, the rankings produced by these two methods are highly concordant at the population level, and we see uniformly high correlations in the ordinal ranks produced by the two methods (Figure 1). In many analyses, the two methods produce virtually identical results when rank is used as a predictor of behavioral or physiological outcomes (Figure 3). In our study system, we have employed and gained insight from both methods (e.g., Levy et al. 2020, Franz et al. 2015). Nonetheless, for simple ordinal rankings, we use the matrix-based approach, with some exceptions (e.g., Levy et al. 2020, Franz et al. 2015), because we see the gains of using the Elo-based ranking method in our study population as modest compared to the benefits of the matrix-based approach.

## Glossary

**Decided agonistic interaction.** Any interaction between two individuals where one individual behaved submissively toward the other and the other gave either neutral or aggressive gestures (see Alberts et al. 2020 for a detailed description). Decided agonistic interactions always have a ‘winner’ and a ‘loser’, but these interactions are not necessarily direct physical fights. An individual might give a submissive gesture (and hence lose a decided agonistic interaction) in response to aggressive behaviors, non-aggressive behaviors, or even no (apparent) behavior at all.

**Diagonal (in a matrix).** The cells that lie on the diagonal that runs from top-left-to-lower-right in a square matrix. The position of an agonistic interaction above or below this line is often helpful for identifying rank rises and falls.

**Loss.** An agonistic interaction in which the individual in question behaved submissively and not aggressively, and its opponent behaved aggressively or neutrally.

**Matrix.** A rectangular array used to show all the agonistic interactions in a group for a specific period of time and age-sex class. A matrix of agonistic interactions is always square. Each column represents a single individual, listed in rank order from left to right (highest-ranked individual at left). Each row likewise represents a single individual, listed in rank order from top to bottom (highest-ranked at top). Each cell contains a number indicating the number of agonistic interactions in which the “column” individual acted submissively to the “row” individual. See the below example, involving individuals ABC, DEF, GHI, and JKL.

|  | ABC | DEF | GHI | JKL |
|---|---|---|---|---|
| ABC |  | 4 | 0 | 2 |
| DEF | 2 |  | 1 | 2 |
| GHI | 0 | 0 |  | 3 |
| JKL | 0 | 0 | 0 |  |

In this example, DEF submitted to ABC in four agonistic interactions, and ABC submitted to DEF in two agonistic interactions. No individuals submitted to JKL. And so on. The top-left-to-lower-right diagonal is empty, because an animal cannot have an agonistic interaction with itself.

Ideally, when the hierarchy is completely linear, the matrix will have 1) only nonzero values above the diagonal, and 2) only zeroes below the diagonal. When it is not possible to order the individuals in such a way as to achieve this, the researcher attempts to minimize the number of agonistic interactions below the diagonal. For example, DEF will not typically be ranked above ABC because if so, the number of agonistic interactions above the diagonal (2) would be higher than the number of interactions below it (4).

**Reversal.** A case in which entries occur below the diagonal of the matrix, i.e., in which an individual has only losses and no wins against an individual ranked below it, or in which both members of a dyad have at least one win against the other. In these cases, members of that dyad will have entries below the diagonal.

**Win.** An agonistic interaction in which the individual in question gave only neutral or aggressive gestures, and its opponent gave only submissive gestures.

## Acknowledgments

We gratefully acknowledge the support of the National Science Foundation and the National Institutes of Health for long-term support for the Amboseli Baboon Research Project, most recently through NSF IOS 1456832, NIH R01AG053308, R01AG053330, R01AG071684, R01HD088558, R01AG075914, and P01AG031719. We thank Duke University, Princeton University and the University of Notre Dame for financial and logistical support. In Kenya, our research was approved by the Kenya Wildlife Service (KWS), the Wildlife Research & Training Institute, the National Environment Management Authority (NEMA), and the National Council for Science, Technology, and Innovation (NACOSTI). We also thank the University of Nairobi, the Institute of Primate Research, the National Museums of Kenya, the members of the Amboseli-Longido pastoralist communities, the Enduimet Wildlife Management Area, Ker & Downey Safaris, Air Kenya, and Safarilink for their cooperation and assistance in the field. Particular thanks go to the Amboseli Baboon Research Project field team (R.S. Mututua, S. Sayialel, J.K. Warutere, I.L. Siodi, G. Marinka, B. Oyath) and camp staff, to T. Wango and V. Oudu for their untiring assistance in Nairobi, and to Jeanne Altmann for her fundamental contributions to the Amboseli baboon research. The baboon project database, BABASE, was designed and programmed by K. Pinc and is expertly managed by N.H. Learn and J.B. Gordon. Joseph Feldblum, Brian Lerch, Chelsea Weibel, Maria Creighton, Maggie Swift, Annie Harshbarger, Laurence Gesquiere, and Clara Howell provided helpful comments on the manuscript. For a complete set of acknowledgments of funding sources, logistical assistance, and data collection and management, please visit http://amboselibaboons.nd.edu/acknowledgements/.

### Appendix: Ethogram of baboon agonistic behaviors and the ABRP procedure for recording decided and undecided agonism interactions

**Table A1.** Table of aggressive behaviors.

| Behavior | Code | Description |
|---|---|---|
| Raised brow or eyelid display | RB | Display of unpigmented skin beneath eyebrows either by raising brow itself or tilting head dorsally; often accompanied by jutting forward of the head. |
| Open mouth face or attempted bite | OF | Jaw held open, teeth not exposed or only slightly exposed, often accompanied by forward jutting of the head. |
| Bobbed head and thorax | BH | Abrupt, rapid raising and lowering of head and trunk; body may show a forward movement component. |
| Ground slapped | GS | Palm of hand or hands struck against ground, often audible. |
| Lunged at | LG | Forward leaping or jumping towards another individual; no lateral movement. |
| Chomped or gave exaggerated chewing motions | CH | Repeated and exaggerated chewing or grinding movements of the jaws, often with extreme lateral excursion, copious salivation and audible grinding. |
| Gave threat yawn or directed gape | GP | Directed gape or yawn-like moving, usually exposing canines. |
| Rubbed muzzle on substrate | RM | Muzzle and chin rubbed laterally on ground or tree limbs. Often accompanied by gaping and chomping. |
| Hit or slapped | HT | Rapid, open-handed striking or attempted striking of another individual. |
| Pushed | PU | Open-handed shoving or attempted shoving of another individual. |
| Grabbed | GB | Closed-handed gripping of another individual, usually brief. |
| Held down | HD | Closed-handed gripping of another individual, forcing that individual against substrate, usually prolonged. |
| Bite | BI | Obvious. |

**Table A2.** Table of submissive behaviors.

| Behavior | Code | Description |
|---|---|---|
| Averted head and/or body | AH | Turned head aside at approach of another individual, often accompanied by fixedly looking away from the other individual, but without the tensing or twisting associated with a cower. |
| Cowered or leaned aside | CW | Lateral flexion of the spine, often from a seated or crouching position; limbs usually held against the body but not always. |
| Gave fear paralysis | FP | Subject leaned forward or lay on ground in rigid, tensed and motionless position. Often accompanied by screaming, urination and defecation. |
| Grimace | GM | Corners of mouth retracted and teeth (often clenched) exposed. Grimaces are sometimes given without being specifically directed toward another individual, e.g., during copulations. Be sure grimace occurs as a directed behavior before you record it as an agonistic interaction. |
| Gave tail-up | TU | Tail held upright, or nearly so, or even 'pointed' forward over back. Note that tail movements occurring during "presents" should not be scored as agonistic behaviors unless they are unambiguously so (in this case there will usually be other submissive gestures also). |
| Gave high-pitched shrill scream or screech | EE | Obvious. |
| Gave cackle or gecker vocalization | IK | Obvious. |
| Repeated rapid glances | GL | Subject glances repeatedly and rapidly in the direction of a conspecific. Sometimes this involves looking over the shoulder or behind; it almost always interrupts the current activity at least briefly. |

**Table A3.** Other agonistic behaviors that are neither clearly aggressive or clearly submissive.

| Behavior | Code | Description |
|---|---|---|
| Walked at or ran at | WT | Obvious. |
| Walked away or ran away | WY | Obvious. |
| Head flagged | HF | Rapid repeated lateral head movements and/or eyelid flashing oriented first toward one individual and then another. We define this behavior as "requesting support"; see Section 10.32. May be accompanied by rapid full or partial closure of the eyelids producing flashes of unpigmented skin. |
| Low-pitched vocalizations such as grunt, roar, bellow and cough | GR | Obvious. |
| Brush past | BP | One animal (usually a male) walks briskly towards a second animal (usually a male) and then suddenly turns and walks away again just when he is very close. Often he literally brushes against the other animal. |

#### Section A1. Procedure for recording decided dyadic agonistic interactions

Decided agonistic interactions between a pair of animals occur when one animal gives submissive gestures (and no aggressive gestures) during the interaction and the other animal gives aggressive or neutral gestures (but no submissive ones). In these cases, the animal that gave the submissive gestures is considered to have “lost” the interaction, and the animal that gave aggressive or neutral gestures is considered to have “won” the interaction. The winner is then generally considered to be higher ranking than the loser (depending on the outcome of other interactions between them that month).

An “aggressive-submissive” decided interaction (or AS interaction) occurs when one animal gives only aggressive gestures and the other animal gives only submissive gestures, or the second animals give a combination of submissive and “other” gestures, within the same continuous agonistic interaction. We consider the animal who gave aggressive gestures to have won the interaction and the animal who gave submissive gestures to have lost the interaction.

An “other-submissive” decided interaction (or OS interaction) occurs when one animal gives only submissive gestures (and sometimes “other” gestures, but no aggressive gestures) and the other animal gives no aggressive or submissive gestures, for instance simply walks towards or past the submitter. We consider the animal who gave no or neutral gestures to have won the interaction and the animal who gave submissive gestures to have lost the interaction.

A displacement (or DS interaction) is also considered a decided interaction. It occurs when neither animal gives aggressive or submissive gestures, but one clearly displaces the other. A displacement occurs in three situations. (1) when one animal moves steadily towards the spot where a second animal is sitting or standing, the second animal moves away when the first is less than five meters away, and the first comes to stand or sit, at least briefly, in the spot where second was; (2) when one animal moves steadily towards the spot where a second animal is sitting or standing, the second moves away when the first is within 5 m, and the first passes right through the spot where the second was; (3) when one animal is on a path that does not intersect with the position of a second animal but comes within one or two meters of it, and the second animal glances in the direction of the first and immediately moves away from its spot. In all these cases, we record that the first animal displaced the second, and we consider the first animal to have won the interaction.

Undecided interactions are those that are dyadic but that do not fall within the criteria for decided encounters; these are often ones in which one or both participants give some mix of aggressive and submissive behaviors. Common undecided interactions include “A-A” (both participants give aggressive gestures), “A-O” (one is aggressive, the other gives some other, non-submissive gesture or ignores the aggressor), and “A-AS” (one animal gives aggressive gestures and the other gives both aggressive and submissive behaviors). “S-S” (both submissive), “AS-AS”, “AS-S” and “AS-O” bouts are also possible but rare.

## Notes

### Competing Interest Statement

The authors have declared no competing interest.

